# The VIL gene *CRAWLING ELEPHANT* controls maturation and differentiation in tomato via polycomb silencing

**DOI:** 10.1101/2021.06.02.446760

**Authors:** Ido Shwartz, Chen Yahav, Neta Kovetz, Alon Israeli, Maya Bar, Matan Levy, Katherine L. Duval, José M. Jiménez-Gómez, Roger B. Deal, Naomi Ori

## Abstract

VERNALIZATION INSENSITIVE 3-LIKE (VIL) proteins are PHD-finger proteins that recruit the repressor complex Polycomb Repressive Complex 2 (PRC2) to the promoters of target genes. Most known VIL targets are flowering repressor genes. Here, we show that the tomato VIL gene *CRAWLING ELEPHANT* (*CREL*) promotes differentiation throughout plant development by facilitating the trimethylation of Histone H3 on lysine 27 (H3K27me3). We identified the *crel* mutant in a screen for suppressors of the simple-leaf phenotype of *entire* (*e*), a mutant in the AUX/IAA gene ENTIRE/SlIAA9, involved in compound-leaf development in tomato. *crel* mutants have increased leaf complexity, and suppress the ectopic blade growth of *e* mutants. In addition, *crel* mutants are late flowering, and have delayed and aberrant stem, root and flower development. Consistent with a role for CREL in recruiting PRC2, *crel* mutants present altered H3K27me3 modifications at a subset of PRC2 targets throughout the genome. Our results uncover a wide role for CREL in plant and organ differentiation in tomato and suggest that CREL is required for targeting PRC2 activity to, and thus silencing, a specific subset of polycomb targets.

**Author summary:** Plants form organs continuously throughout their lives, and the number and shape of their organs is determined in a flexible manner according to the internal and external circumstances. Alongside this flexibility, plants maintain basic developmental programs to ensure proper functioning. Among the ways by which plants achieve flexible development is by tuning the pace of their maturation and differentiation, at both the plant and organ levels. One of the ways plants regulate the rate of maturation and differentiation is by changing gene expression. Here, we identified a gene that promotes plant and organ maturation and differentiation. This gene, *CRAWLING ELEPHANT* (*CREL*) acts by bringing a repressing complex to target genes. We show the importance of CREL in multiple developmental processes and in the expression of multiple genes throughout the tomato genome.

## Introduction

Polycomb Repressive Complex 2 (PRC2) is a conserved complex that represses gene expression by trimethylating lysine 27 of histone H3 proteins (H3K27me3)[1–3]. PRC2 activity counteracts, and is counteracted by, the transcription-promoting functions of trithorax-group proteins [4]. The core PRC2 is composed of 4 subunits. In plants, some of these subunits are encoded by small gene families, allowing the formation of multiple, distinct complexes. Different plant PRC2 complexes have been shown to regulate specific developmental processes such as endosperm development, flowering time and flower development [2,3]. As PRC2 complexes do not have DNA binding domains, they are recruited to target loci by interacting proteins [2,5–7]. One of the most characterized PRC2-regulated processes in Arabidopsis is the induction of flowering in response to prolonged cold, termed vernalization. In response to vernalization, PRC2 promotes flowering by silencing the flowering inhibitor FLC. The vernalization-specific VRN-PRC2 complex is recruited to FLC by complexing with PHD proteins from the VERNALIZATION INSENSITIVE 3-LIKE (VIL) family [7–10]. In Arabidopsis, the VIL family consists of 4 members, including VIN3 and VRN5. Vernalization induces VIN3 expression, while VRN5 is expressed constitutively. VIL proteins also repress additional members of the FLC family during vernalization, and VRN5 and VIL2 are also involved in other flowering pathways [7,8,11,12].

VIL proteins have been identified from several plant species [13–22]. They have been shown to promote flowering in all tested species, including species that do not have an FLC ortholog and/or do not respond to vernalization. In rice, the *OsLF* and *OsLFL1* genes encode transcription factors that inhibit flowering and have been identified as VIL targets [15,17]. A VIN3 ortholog has also been identified in tomato [21]. While the vast majority of research on VIL proteins concerned their involvement in flowering induction, several studies reported additional developmental effects. For example, Arabidopsis *vrn5* mutants had increased leaf curling, increased numbers of petals, and distorted siliques [10]. In rice, *osvil3/leaf inclination2* (*lc2*) mutants had an altered leaf angle, curled leaves and severe sterility, and OsVIL2 was found to affect spikelet development, branching and grain yield [13,14,16,23]. Silencing the *Brachypodium distachyon BdVIL4*, which is similar to *VIN3*, led to increased branching [18]. Pepper *cavil1* mutants affect leaf development, apical dominance and branching [22]. However, the knowledge about the involvement of VIL proteins in these and other developmental processes is limited, and their role in compound-leaf development has not been explored. In addition, it is not clear whether VIL proteins recruit PRC2 mainly to targets involved in the induction to flowering or whether they have broader roles in plant development.

Tomato plants have compound leaves, which are composed of multiple leaflets [24]. The elaboration of compound leaves depends on slow maturation of the developing leaf, which enables an extended organogenesis activity at the leaf margin, during which leaflets are formed [25–29]. Leaflets are formed by differential growth at the leaf margin, where regions of blade growth are separated by intercalary regions of growth inhibition [30]. Auxin has been shown to promote growth and its response is inhibited in the intercalary domains [31–37]. Mutations in the tomato gene *SlIAA9/ENTIRE* (*E*), encoding an auxin-response inhibitor from the Aux/IAA family that specifies the intercalary domain, result in simplified leaves due to ectopic blade growth in the intercalary domain [31,38,39].

Here, a screen for suppressors of the *e* simple-leaf phenotype identified the *crawling elephant* (*crel*) mutant, which substantially suppresses the ectopic blade growth of *e*. We found *CREL* to encode a tomato VIL gene, related to Arabidopsis *VIL1/VRN5. crel* mutants affect many aspects of tomato development, including plant and organ maturation. Comparison of H3K27me3 modifications between wild type and *crel* plants showed that CREL affects these modifications at some PRC2 targets and not others. Therefore, CREL promotes maturation throughout the plant life by promoting selective deposition of H3K27me3 and gene silencing at a subset of PRC2 targets.

## Results

### *crawling elephant* (*crel*) mutants suppress *entire* (*e*) and have very compound leaves

*entire* (*e*) mutants, mutated in a tomato Aux/IAA gene, have simplified leaves in comparison to the wild-type compound leaves [31,37,39,40] (Fig 1A, B). To identify genes that are involved in compound-leaf development, we generated an Ethyl Methane Sulfonate (EMS) mutant population in the background of *e*, and screened for suppressors of the *e* simplified leaf phenotype. This screen identified the *crawling elephant-1* (*crel-1*) mutant as a strong *e* suppressor. *e crel-1* double mutants had distinct, clearly separate primary leaflets, and occasionally had secondary leaflets, in contrast to the mostly entire leaf shape of single *e* mutants (Fig 1A-C). To characterize the unique *crel-1* phenotype, we backcrossed *crel-1* to the parental line (*Solanum lycopersicum* M82), and identified single *crel-1* F2 individuals (Fig 1D). Leaves of single *crel-1* mutants were much more compound than wild-type leaves, with a similar number of primary leaflets but many more secondary leaflets than the wild type. In contrast to wild type-leaves, *crel-1* leaves also had tertiary leaflets (Fig 1A, D, J, K). Therefore, *crel* mutants suppress the *e* simplified-leaf phenotype, and forms many more leaflets in both the wild type and the *e* backgrounds. Interestingly, previously identified *e* suppressors such as *slmp* and *slarf19a,b* had a reduced number of leaflets in the respective single mutants [36].

**Fig 1.**
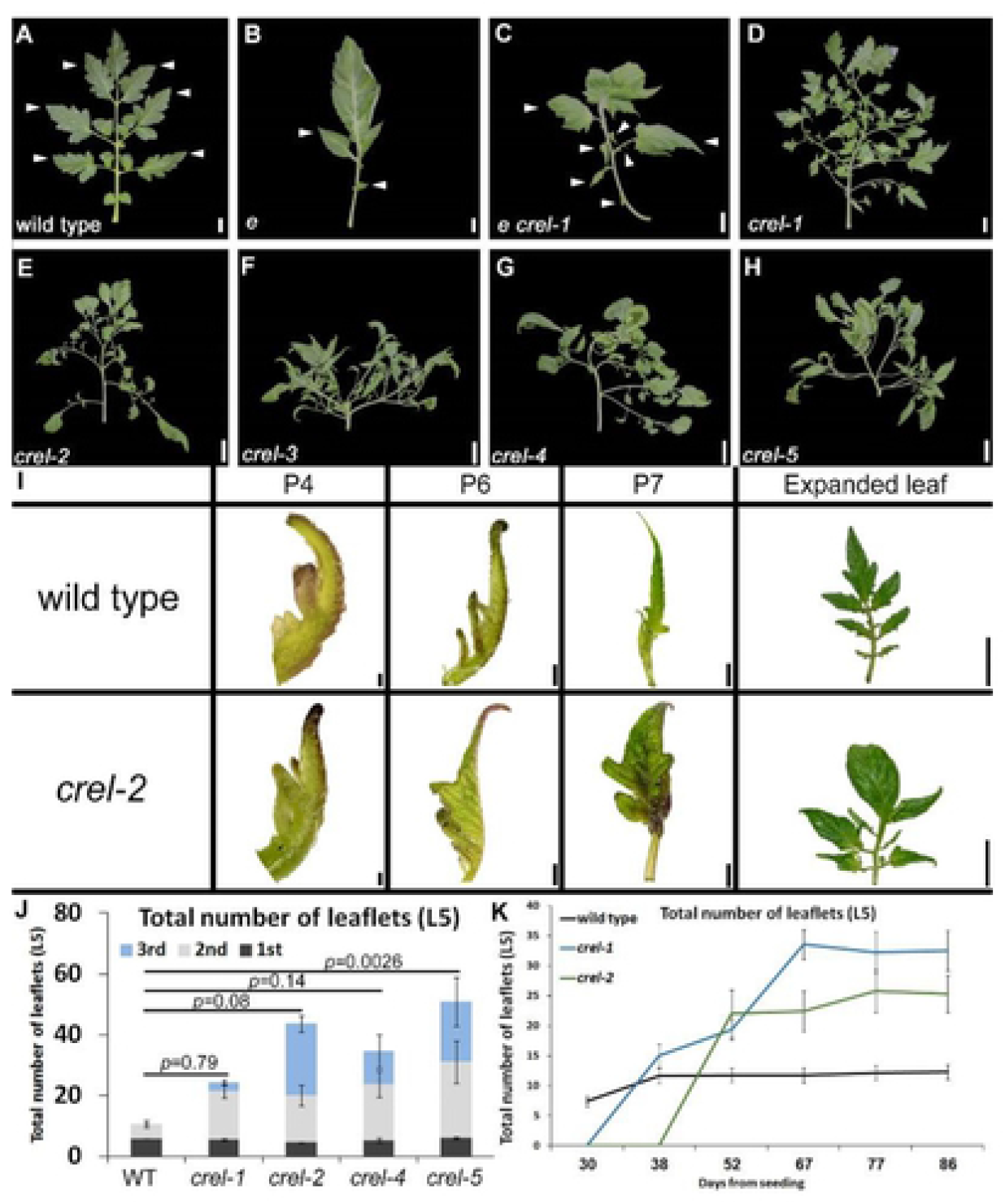
Mutating *crel* suppresses the e simple leaf phenotype and produces very compound leaves. (A-H) Mature 5^th^ leaves of the indicated genotypes. A-C suppression of *e* by *crel-1*; D-H: leaves of 5 different *crel* alleles. Scale bars: 2cm. (I) Early leaf development in the fifth leaf of wild type and *crel-2*. P4-P7 designate the developmental stages, where P4 is the forth youngest leaf primordium. Scale bars: 0.1 mm (P4), 0.5 mm (P6), 2 mm (P7), 2 cm (expanded leaf). (J) Quantification of the number of leaflets in a mature 5^th^ leaf of the indicated *crel* alleles, compared to the wild type. 1^st^, 2nd and 3^rd^ represent primary, secondary and tertiary leaflets, respectivley, where primary leaflets arise from the rachis, secondary leaflets arise from primary leaflets etc. (K) Leaflet production over time by the fifth leaf (L5) of the indicated genotypes.

We identified several additional *crel* alleles from the Menda EMS and fast neutron mutant population [41], in the M82 background, and confirmed allelism by complementation tests. The alleles showed a range of phenotypic severities, including a diverse increase in leaflet number (Fig 1E-H, J, K). Similar to *crel-1, crel-2* also suppressed the *e* simplified leaf phenotype (S1 Fig A-D).

### CREL acts during relatively late stages of leaf development

To investigate the timing of the effect of *crel* mutants on leaf development, we compared leaf development between wild type and *crel-1* plants. Early stages of leaf development were very similar between wild type and *crel-1* plants when similar developmental stages were compared, although the terminal leaflet expanded earlier in *crel-1* (Fig 1I). However, the rate of leaf and leaflet initiation was much slower in *crel-1* mutants than in the wild type (S1 Fig E). At later stages of leaf development, when wild type leaves stopped generating new leaflets, *crel-1* and *crel-2* leaves continued to form leaflets more than a month later (Fig 1K). Therefore, *crel* leaves develop slower than the wild type, and while the terminal leaflet appears to differentiate early, overall leaf differentiation is substantially delayed in *crel* mutants.

To further characterize this effect of *crel* on leaf development, and understand the timing and developmental context of CREL action, we analyzed its genetic interaction with mutants that affect the developmental window of leaflet morphogenesis. Leaflets are formed during the morphogenesis stage of leaf development, which follows leaf initiation and precedes leaf expansion and differentiation [24,25,42,43]. The elaboration of compound leaves depends on an extended morphogenesis stage. The CIN-TCP transcription factor LANCEOLATE (LA) and the MYB transcription factor CLAUSA (CLAU) promote maturation and differentiation and thereby restrict the morphogenetic window [26,44–47]. *La-2* is a semi-dominant mutant in which *LA* is expressed precociously due to a mutation in the miR319 binding site. This accelerates leaf differentiation and leads to a simple leaf form (Fig 2A, C). *La-2* was epistatic to *crel-1* (Fig 2 B-D), indicating that the morphogenetic window in *La-2* is terminated before the timing of CREL action, in agreement with the relatively early effect of *LA* and late effect of *CREL* on leaf development. Leaves of loss-of-function *clau* mutants have an extended morphogenetic window, leading to a substantial increase in leaf complexity and leaflet number (Fig 2E) [45,48]. *crel-1 clau* double mutants had very complex leaves (Fig 2E, F), suggesting that CREL acts in parallel with CLAU to restrict leaf elaboration and promote maturation. Removing the activities of both regulators leads to prolonged, extensive leaflet morphogenesis. Together, these results suggest that CREL acts in relatively late stages of leaf development to promote maturation and differentiation.

**Fig 2.**
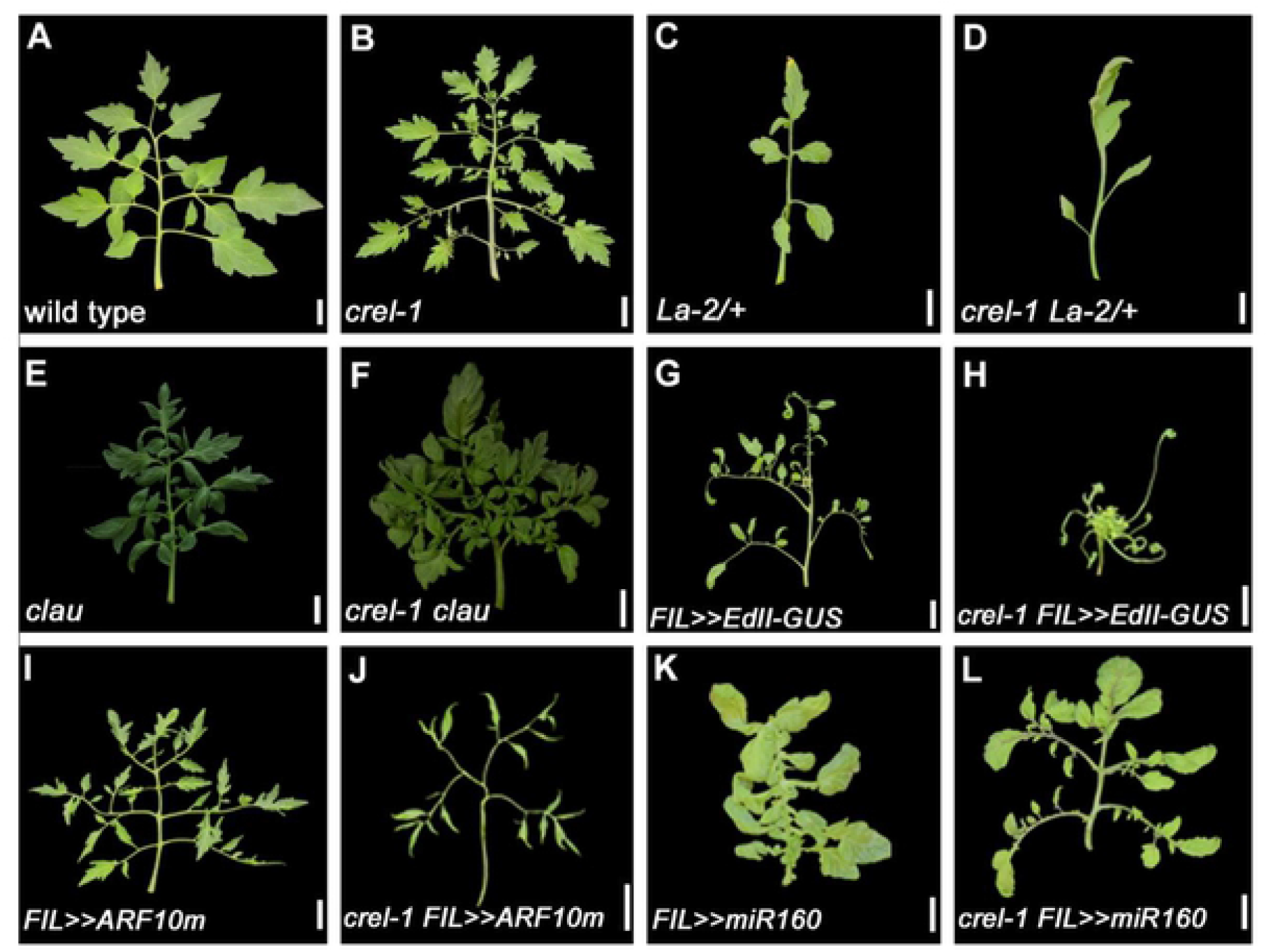
CREL acts relatively late in leaf development to promote differentiation and blade growth. (A-F, I, J) Mature 5^th^ leaves. (K,L) Mature 9^th^ leaves. (G, H) whole plants. Scale bars: 2cm. *FIL*>>gene refers to genotypes generated by the LhG4-OP transactivation system, where the gene is expressed in the *FIL* expression domain. The *FIL* promoter is expressed in leaf primordia [1]; *EdII-GUS* is a stabilized form of E fused to the GUS reporter; *ARF10m* is a mutant form of ARF10 that is mutated in the miR160 binding site. A-D - epistasis of *La*-2/+, in which differentiation is accelerated, over *crel*. E, F - enhancment of crel by *clau*, in which differentiation is delayed. G-J – enhancment of genotypes with narrow leaves due to reduced auxin response by crel. K, L – suppression of the ectopic blade growth of *FIL*>>*miR160* by crel, similar to the effect on *e*.

The suppression of the *e* phenotype by *crel* raised the question of whether CREL is also involved in the differential growth at the leaf margin that leads to the formation of separate leaflets. To address this question, we crossed *crel* to mutants affected in auxin-mediated blade growth. Ectopic expression of a stabilized form of E resulting from a mutation in domain II of the E (IAA9) protein (EdII) resulted in leaflet narrowing [39], (Fig 2G). This effect was strongly enhanced in the *crel-1* background (Fig 2H), suggesting that E inhibits and CREL promotes blade expansion, but they act in at least partially parallel pathways. Similarly, *crel-1* enhanced the narrow blade phenotype resulting from ectopic expression of a miR160-resistant ARF10, a negative regulator of blade expansion (*FIL*>>*ARF10m*, Fig 2I, J). In agreement, *crel-1* suppressed the ectopic blade growth resulting from ectopic expression of miR160, which negatively regulates ARF10 and 4 additional ARF proteins (*FIL*>>*miR160*, Fig 2K, L) [37]. The suppression of *FIL*>>*miR160* was more prominent in later leaves than in early leaves. Interestingly, leaflet number was reduced in *FIL>>ARF10 crel-1* relative to both single mutants, and *crel-1 FIL>>EdII-GUS* plants were extremely small with almost no leaflets, suggesting that extreme repression of lamina growth leads to a reduction in leaflet formation and overall growth. Together, these results suggest that CREL promotes blade growth during compound-leaf development, and acts at least partially in parallel to auxin.

### CREL is a VRN5 homolog

To identify the *CREL* gene, we genetically mapped the *crel-1* mutation using an F2 mapping population from a cross between the *crel-1* mutant, in the *Solanum lycopersicum* M82 background, and *S. pimpinellifolium. crel-1* was mapped to the short arm of chromosome 5. Further mapping was hampered by an introgression of *S. pimpinellifolium* sequences in the M82 line in this region [49]. We therefore used RNAseq to identify possible causative mutations in *crel-1* and *crel-2*, which led to the identification of mutations in the gene *Solyc05g018390* in both *crel-1* and *crel-2*. In *crel-1*, a G to A substitution at position 4264 from the transcription start site (TSS) led to a stop codon in exon III. The fast neutron allele *crel-2* contains a 12,826-bp-long deletion, which results in the elimination of exon I and II and part of exon III (Fig 3A). Sequencing the Solyc05g018390 gene in two additional *crel* alleles identified a 1-bp deletion in the first exon at position 322 from the TSS in *crel-3*, and an A to T substitution in position 3630 leading to a stop codon in the third exon in *crel-5* (Fig 3A). We therefore concluded that Solyc05g018390 is *CREL. CREL* is predicted to encode a plant homeodomain (PHD) finger protein (Fig 3A, B). It is most similar to the Arabidopsis VRN5 gene.

**Fig 3.**
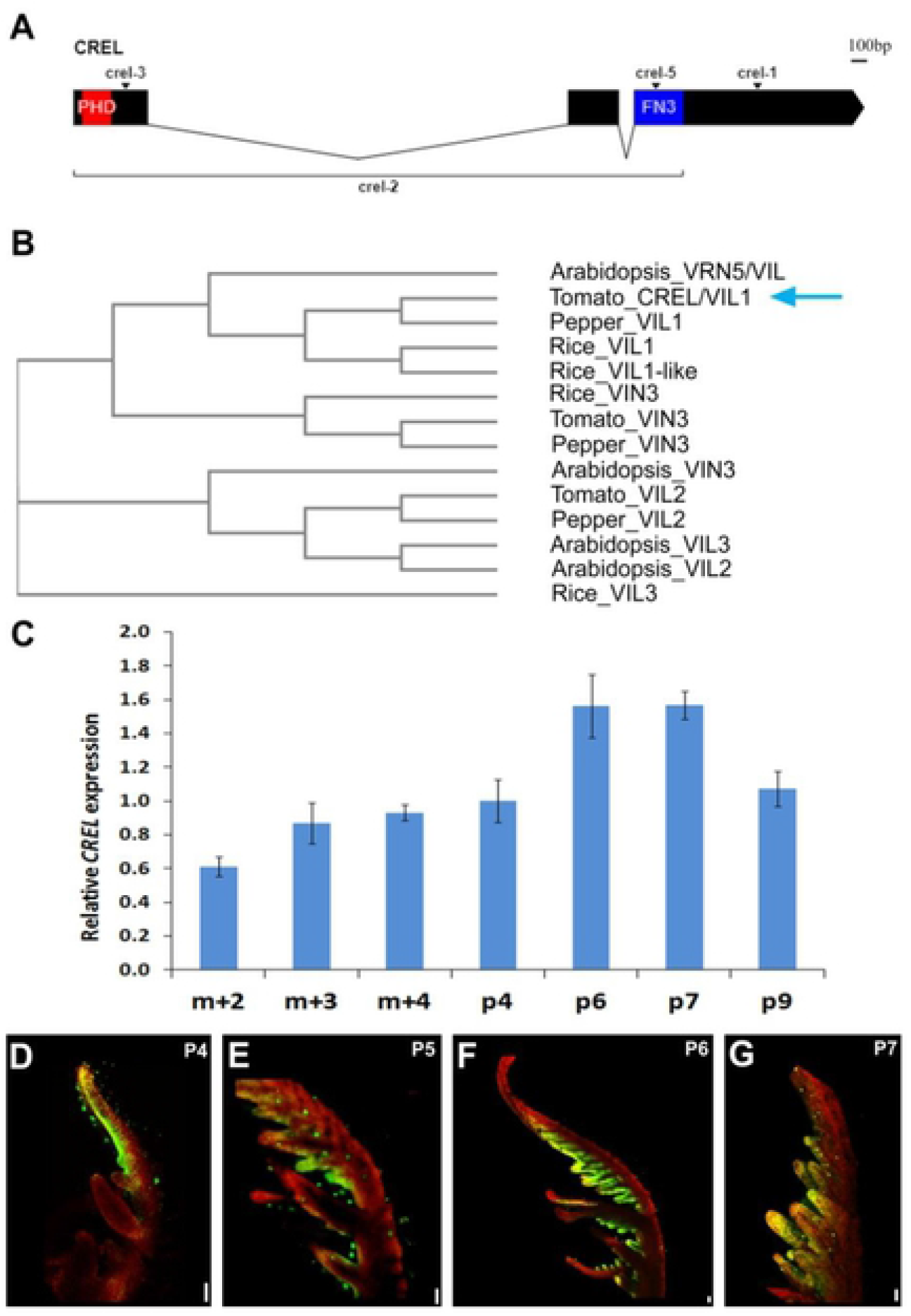
CREL encodes a VRN5/VIL1 homolog expressed at late stages of leaf development. (A) A diagram of the CREL (Solyc05g018390) gene. The boxes indicate exons and the combining lines introns. The location of the mutation in 4 *crel* alleles is indicated. (B) A phylogenetic tree of the tomato, Arabidopsis, rice and pepper VIL proteins, constructed using Clustal Omega (https://www.ebi.ac.uk/Tools/msa/clustalo/). The blue arrow points to CREL. (C) qRT-PCR analysis of CREL mRNA expression at successive developmental stages of the 5th leaf. m+2, 3, or 4 represents the meristem and the 2, 3, or 4 youngest leaf primordia, respectively. P4-P9 represent isolated leaf primordia at the respective developmental stage (see figure 1). Error bars represent the SE of at least three biological replicates. (D-G) Confocal images of leaf primordia of the indicated stages, expressing *pCREL*>> *YFP*, using the transactivation system, as in figure 2. P4-P7 represent the 4^th^-7^th^ youngest leaf primordia, respectively. In G, a leaflet from a P7 primordium is shown. Scale bars: 0.1 mm.

### CREL is expressed in expanding blades

We characterized the expression of *CREL* in the fifth leaf produced by the plant, at different developmental stages, to examine how its expression correlates with its activity. *CREL* was expressed throughout leaf development, with relatively low expression in apices containing the SAM and very young P1-P3 primordia. Later, its expression was gradually upregulated, peaking at P6/P7 (Fig 3C). To spatially localize *CREL* in leaf primordia, we cloned a 2960-long *CREL* promoter and used it to generate a CREL driver line in the transactivation system [50,51]. In developing leaves, the *CREL* promoter drove expression in leaf margins. In agreement with the qPCR experiment, expression appeared lower in young primordia, and increased from P4 on. Expression was mainly visible in expanding regions of the leaf margin, the terminal leaflet at P4, and the expanding leaflets at P6 and on (Fig 3D-G). The expression appeared to follow the basipetal differentiation wave of the leaf, with strong expression first appearing in the terminal leaflet, which is the first to expand and differentiate, and then progressing basipetally in expanding leaflets. This leaf expression pattern is compatible with the *crel* leaf phenotype, which starts to differ from the wild type around the P5 stage (Fig 1I), and with the genetic interactions showing that *crel* affects leaf maturation and blade formation (Fig 2).

### *CREL* promotes multiple aspects of plant maturation and differentiation

In addition to their effect on leaf differentiation and patterning, additional differentiation processes were also delayed and/or impaired in *crel* mutants. *crel* plants failed to maintain an upright position and the plants exhibited a sprawling growth habit. Similar to the effect on leaf shape, this phenotype developed at a relatively late stage of plant development (Fig 4A, B). To further understand the role of CREL in plant development, we overexpressed the *CREL* gene under the control of the 35S promoter. *CREL* mRNA expression increased only slightly in *35S:CREL* plants (S2 Fig A), and the phenotypic effect was subtle (Fig 4A, B), but when mature plants were compared, *35S:CREL* plants were slightly taller than *crel-2* mutants (S2 Fig B). To understand the basis for the “crawling” phenotype, we dissected developing wild-type and *crel-2* stems at successive developmental stages. We sectioned the internode between the cotyledons and the first leaf from different plants grown together, between the ages of 3 and 10 weeks. In three-week-old plants, *crel-2* stems were narrower than the wild type with nearly normal although slightly less developed vascular bundles (Fig 4C, D, J, K). *crel-2* vasculature continued to develop slower than the wild type, and ceased maturation and differentiation prematurely. This resulted in a thin and undeveloped xylem in *crel-2* stems. Specifically, *crel-2* stems failed to complete a vascular cylinder, and had only partial secondary xylem development (Fig 4E-H, L-O). The reduction in supporting tissue likely contributes to the reduced strength of the *crel* stem. Stem sections of 10-week-old *35S:CREL* plants were similar to those of the wild type but appeared to mature more slowly (Fig 4I, P).

**Fig 4.**
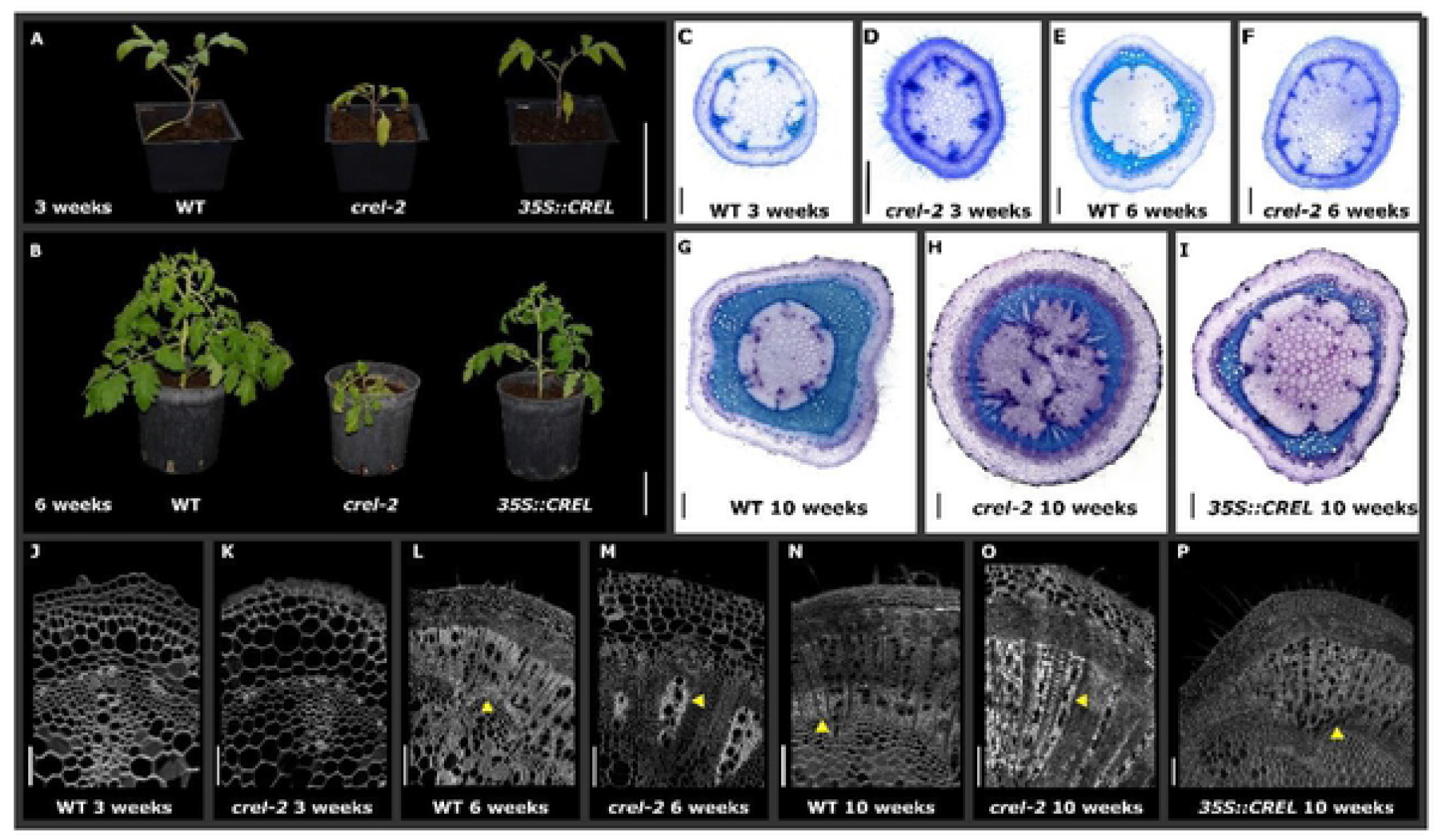
CREL promotes stem vasculature maturation and differentiation. (A, B) Whole plants of the indicated genotypes and ages. Scale bars: 10 cm. (C-I) stem cross sections, dissected from the first internode of the plant (between the hypocotyl and first leaves) at the indicated times after germination and stained with Toluidine blue. Scale bars: 500 mm. (J-P) Confocal images of stem cross sections, taken from the first internode of the plant at the indicated times after germination. Yellow arrowheads point to differentiated (WT) or undifferentiated (*crel-2*) xylem/ vasculature. Scale bars: 100 mm (J,K); 200 mm (M); 500 mm (L,N-P).

Root vasculature development was also delayed and impaired in *crel* mutants. *crel-2* roots were narrower than wild-type roots, and their vascular tissue developed slowly and failed to reach full differentiation (Fig 5A-F). To investigate the effect of *crel* on the root system as a whole, wild type and *crel-2* plants were grown hydroponically, and root volume and length were calculated at successive times. Root volume and length were reduced in *crel-2* plants, and the difference increased with time, although the difference was statistically significant at one of the time points only (Fig 5G,H). Therefore, CREL plays an important role in root development and differentiation.

**Fig 5.**
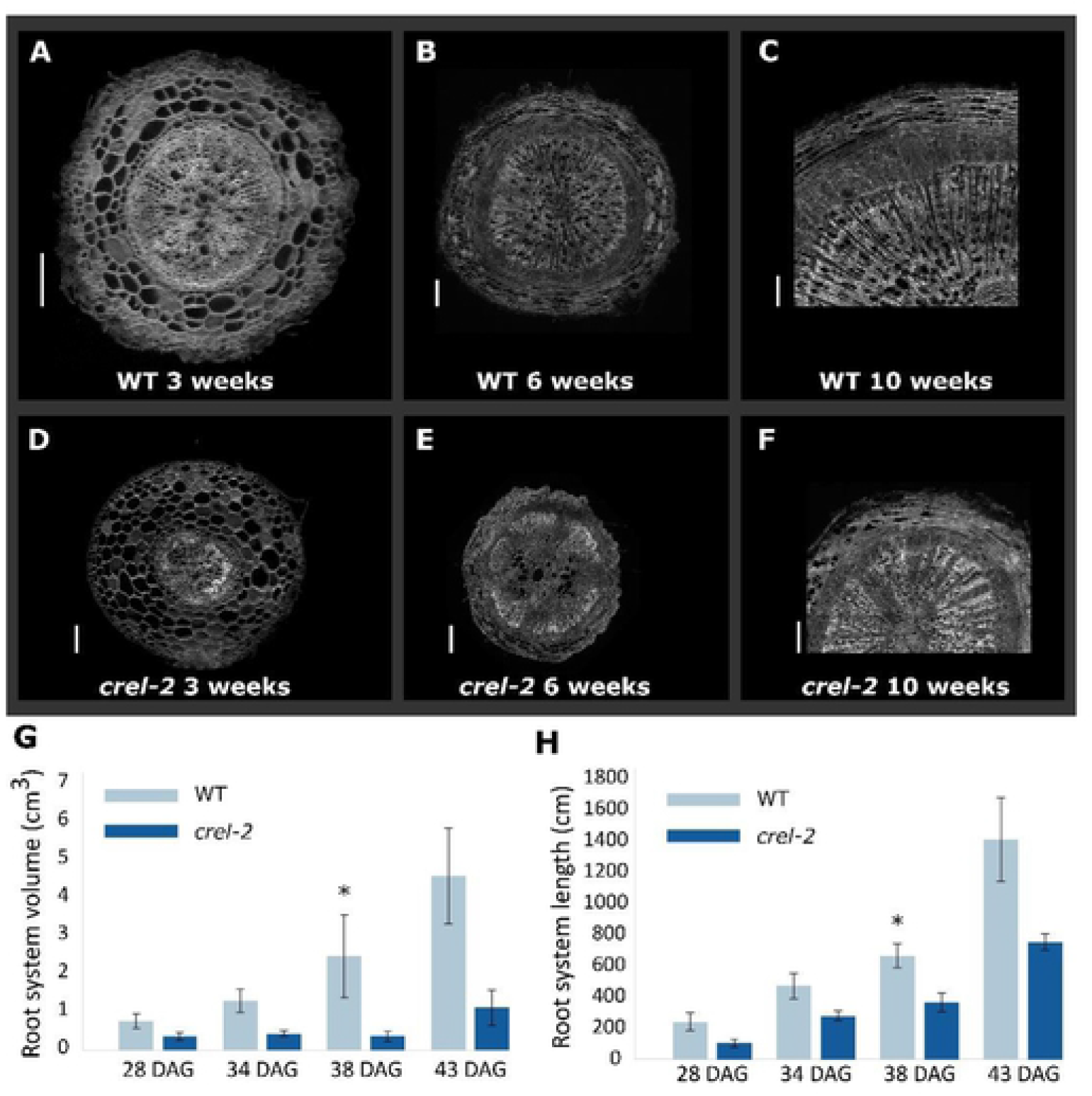
CREL promotes root development and differentiation. (A-F) Confocal images of root cross sections of the indicated genotypes dissected from the upper part of the primary root. Scale bars: 200μm (A, B, G); 500μm (C-F). (G) Root system volume at different times after germination, calculated with WinRhizo software. Shown are averages and SE of 3 plants (n=3). Asterisks indicate statistically significant differences between *crel-2* and WT, by Student’s t test, *p < 0.05. (H) Root system length at different times after germination (DAG), calculated withWinRhizo software. Shown are averages and SE of 3 plants (n=3). Asterisks indicate statistically significant differences between *crel-2* and WT, by Student’s t test, *p < 0.05.

The *crel* mutation also affected flowering time and flower development. *crel-1* and *crel-2* mutants flowered much later than the wild type, after producing 12-13 leaves, compared to 6 leaves in the wild type (Fig 6A). *35S:CREL* plants flowered slightly earlier than the wild type, but this effect was statistically significant in only one of the lines (S2 Fig C). Mature *crel* flowers were not fully developed, had short and distorted organs and were sterile (Fig 6B, C). Early flower development was similar between wild type and *crel* plants, except for the sepals that were curled backwards in *crel*, resulting in an open bud where the inner organs were not covered by the sepals. However, *crel* flower organs ceased development and growth prematurely (Fig 6D-G). *35S:CREL* flower development was similar to that of the wild type and the plants were fertile (data not shown). Therefore, *crel* mutants were delayed in multiple developmental pathways. In some cases such as flower, stem, and root development, these organs failed to properly differentiate, while in others, such as leaf development and flowering time, they differentiated substantially slower than the wild type. Overall, *crel* plants had aberrant plant and organ structure, which resulted in weak and sterile plants.

**Fig 6.**
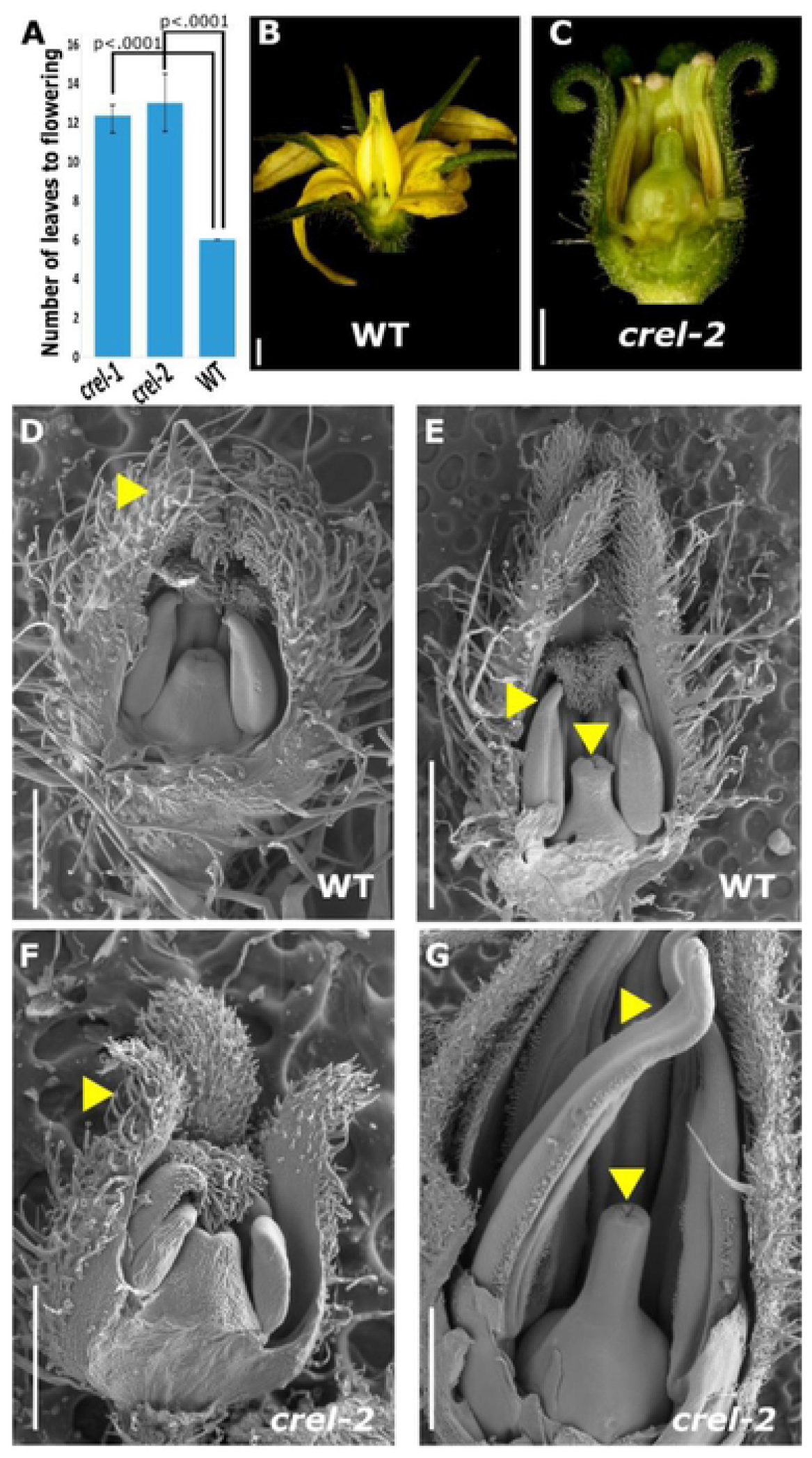
CREL promotes flowering, floral organ growth, and differentiation. (A) Flowering time, measured by number of leaves produced before flowering, of the indicated genotypes. Error bars represent the SE; p-values indicate differences from WT, as determined by Dunnett’s test. n=12 (wt, *crel-1*) and 4 (*crel-2*). (B, C) Stereoscope images of mature flowers. Scale bars: 1mm. (D-G) Scanning electron microscope (SEM) micrographs of the indicated genotypes at 2 early developmental stages. D, F - 0.5 mm long stage 6 flowers; E, G - 1 mm long stage 11 flowers (according to [2]. Yellow arrowheads point to normal (WT, D) and distorted (*crel-2*, F) young petals, and to normal (WT, E) and distorted (*crel-2*, F) stamens and stigma. Scale bars: 1mm (E, G); 0.5mm (D, F).

### CREL mediates H3K27me3 modifications at a subset of polycomb-silenced genes

Homology of *CREL* to the Arabidopsis *VRN5* gene suggested that it may be involved in the repression of gene expression by promoting PRC2-mediated H3K27me3 modification. To test this prediction, we performed ChIP-seq for the H3K27me3 modification in shoot apices of 4-week-old wild-type and *crel-2* plants (Supplemental Table 1). In wild-type tissue, H3K27me3 was found to be significantly enriched at a total of 13,849 sites, mostly over gene bodies, as expected. In the *crel-2* mutant, H3K27me3 appeared to be completely lost at 6,762 of these sites (48.8%) normally enriched with H3K27me3 (Fig 7A), supporting the hypothesis that CREL normally guides deposition of H3K27me3 at a subset of PRC2 target genes. Interestingly, 4,789 sites actually show significant increases in H3K27me3 in the *crel-2* mutant (Fig 7A). The vast majority of these sites are normally enriched for H3K27me3 in WT, suggesting that in the absence of CREL, excess PRC2 activity is directed to the remaining target genes.

**Fig 7.**
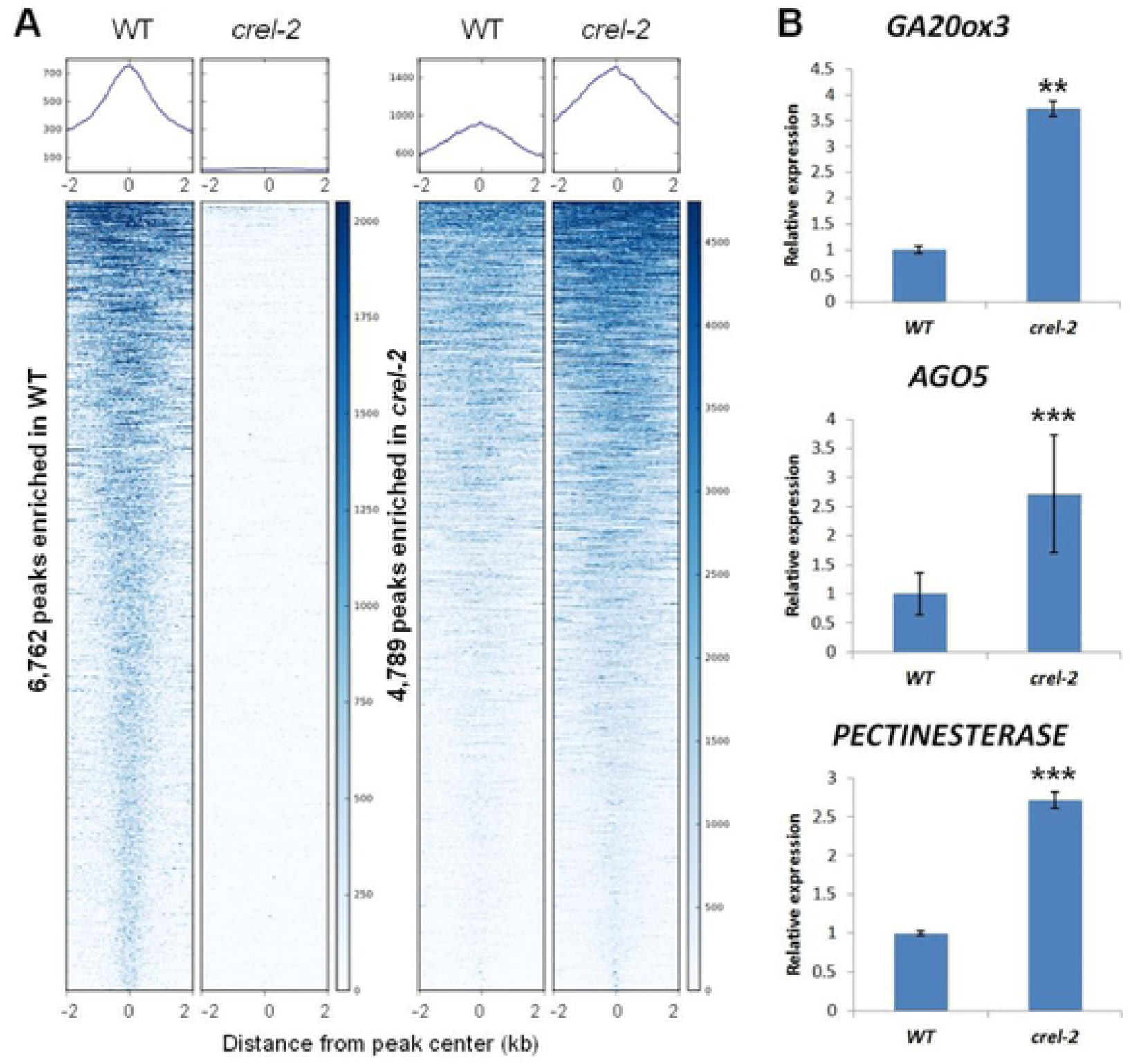
CREL mediates H3K27me3 modifications at a subset of polycomb-silenced genes. (A) Average plots and heatmaps show H3K27me3 enrichment in shoot apices of WT and *crel-2* plants. The left pair of panels show the 6,762 H3K27me3-enriched sites in WT (out of a total of 13,849 sites) where the modification is significantly depleted in *crel-2* mutants. The right pair of panels show 4,789 H3K27me3-enriched sites where the modification level is higher in the *crel-2* mutant. The majority of these sites are normally enriched with H3K27me3 in WT and the levels become higher in *crel-2*. (B) qRT-PCR analysis comparing the mRNA expression of *GA20oxidase 3* (*GA20ox3*, Solyc11g072310), *ARGONAUTE 5* (*AGO5*, Solyc06g074730) and *PECTINESTERASE* (Solyc02g080200) in primordia of the fifth leaf produced by the plant at the P5 stage from wild type (WT) and *crel-2* plants. These were among the genes in which H3K27me3 was lost in *crel-2* in comparison to the wild type. The bars represent the average of 3-5 biological replicates, and error bars indicate SE. Asterisks indicate statiscally significant diffrences, determined by students t-test, **P<0.01, ***P<0.001.

To examine the effects of these H3K27me3 re-distributions on target gene expression, we used reverse transcription followed by quantitative PCR (RT-qPCR) to examine the levels of several target transcripts in young leaf primordia. The tested genes were among the genes that were normally enriched for H3K27me3 in WT and lost modification in *crel-2*, and were upregulated in *crel* mutants in the RNAseq used for the identification of the CREL gene. We compared the expression of three such genes, *GA20oxidase 3* (*GA20ox3*, Solyc11g072310), *ARGONAUTE 5* (*AGO5*, Solyc06g074730) and *PECTINESTERASE* (Solyc02g080200), in P5 primordia of the fifth leaves produced by the plant. As expected, these genes showed increased transcript levels in the mutant (Fig 7B). These results are all consistent with the known role of H3K27me3 in gene silencing and further support the validity of our ChIP-seq data.

## Discussion

VIL proteins have been shown to affect flowering in several plant species, by repressing the expression of flowering repressors, such as *FLC* in Arabidopsis. In addition to their effect on flowering, VIL genes were found to affect an array of developmental processes in different species. This work identifies the tomato VIL gene *CREL* as a mediator of diverse developmental processes, via the modulation of H3K27me3 modifications in many genes throughout the tomato genome, likely by recruiting PRC2 complexes to a subset of their target genes.

### CREL promotes plant and organ maturation

So far, VIL genes have been mainly implicated in flowering time [7,11,13,15,17–19,22,52]. Here, we uncover a much broader role for this gene family in plant development, as revealed from the phenotypes and the effect on H3K27me3 modification. *crel* mutants are affected in many aspects of plant maturation and differentiation in addition to the delay in flowering time. *crel* mutants flower late and have delayed leaf maturation, resulting in an extended leaf morphogenesis and more compound leaves. Interestingly, while flowering and leaf maturation eventually occur in *crel*, stem, root, and flower differentiation are impaired in *crel* and these organs do not reach full differentiation and function. CREL accumulates relatively late during leaf development, thus enabling prolonged morphogenesis. Recently, a growth-rate dependent mechanism of controlling VIN3 accumulation in the cold has been described [53]. It would be interesting to understand the mechanism by which CREL expression is delayed during organ maturation to enable timed maturation and differentiation.

Other genes involved in the induction of flowering were also shown to affect maturation and differentiation in additional developmental aspects. For example, the tomato flowering inducer SFT, the ortholog of FT, was shown to promote leaf maturation and affect stem differentiation [54]. Recently, SFT was shown to specifically affect secondary cell wall biosynthesis in tomato stems [55]. FT was also proposed to promote maturation and termination in additional species [56,57]. The pepper *cavil1* mutants, impaired in the pepper *CREL* ortholog, have reduced vascular development but wider stems [22]. Therefore, both precocious and delayed differentiation impairs the final form and function of stems. In *Cardamine hirsuta*, plant maturation and flowering was shown to be coordinated with age-dependent changes in leaf shape in plants with variable FLC activity [58]. In contrast to CREL and SFT, which promoted all aspects of plant and organ maturation, tomato CIN-TCPs were shown to promote leaf maturation but delay plant maturation, while AP1/FUL MADs BOX genes promoted plant maturation and delayed leaf maturation [59,60]. Overall, similar to *CREL*, genes that have been implicated mainly in the promotion of flowering in Arabidopsis were found to promote a wide range of differentiation and maturation aspects.

The involvement of CREL in plant and organ differentiation is in agreement with a role in mediating PRC2 activity. A common function of PRC2 genes in plants is the maintenance of a differentiated state, and *prc2* mutants in both mosses and seed plants have phenotypes related to dedifferentiation and overproliferation [1,2,61]. Therefore, CREL may aid in recruiting PRC2 to differentiation-related target genes.

VIL genes from other species have also been shown to affect other developmental processes in addition to flowering. [10,13,16,18,22,23]. Interestingly, beside the common effect on flowering time, the specific developmental effects only very partially overlap among these species. This suggests that the VIL family may be used as a tool for developmental innovations, recruiting an existing tool to different processes. Specifically interesting in this respect is the comparison between tomato and pepper, which are closely related species that differ in several key developmental features, such as flowering architecture and leaf shape. *cavil1* mutants have reduced vasculature development, increased plant and organ size, increased branching and reduced angle of axillary branches [22]. Interestingly, only some of these additional phenotypes overlap with *crel*. In contrast to the simple leaves of pepper, tomato leaves are compound, with several orders of leaflets. This is correlated with faster differentiation of the pepper leaf, similar to tomato *La-2/+* mutants [26]. The current work revealed an important role for CREL in the development of the compound leaf, with an effect on both the rate of differentiation and leaf patterning (Fig 1 and 2), further supporting the notion that VIL genes have been recruited to diverse, partially species-specific processes.

We propose that, in addition to its general effect on growth and differentiation, CREL also promotes blade growth in developing tomato leaves. *crel* mutants suppress the ectopic blade outgrowth of *e* mutants and miR160-overexpressing plants (Fig 2K, L). Furthermore, *crel* enhances the narrow-leaf phenotype caused by overexpression of *E* or miR160-targeted ARFs. In addition, *CREL* expression is elevated in later stages of leaf development when the blade begins to expand, and the *CREL* promoter shows high expression in growing regions of the leaf margin (Fig 3D-G)). The genetic interactions between *crel* and auxin-related mutants suggest these auxin mediators and CREL act via independent pathways to regulate blade growth. Therefore, CREL likely promotes blade growth either downstream of auxin or through an at least partially parallel pathway. As most effectors of compound leaf development have been shown to affect either the organ-level differentiation rate (for example LA and CLAU) or local differential growth (E, CUC, SlMP) [24], it is interesting that CREL appears to affect both aspects.

### CREL affects H3K27me3 modifications throughout the tomato genome

Only a handful of VIL targets have been identified so far, most of which are related to their role in promoting flowering. In Arabidopsis, *FLC* and FLC-related genes are targeted by different VIL protein in specific flowering pathways [8,11]. In rice, the flowering inhibitors *OsLF* and *OsLFL1* were identified as VIL targets [15,19,62]. In addition, a cytokinin catabolism gene from the CKX family and the bud-outgrowth inhibitor OsTB1 have been identified as a VIL target in rice [16,23]. The microRNA miR156 was proposed as a target of BdVIL4 in *Brachypodium* [18], and several putative targets have been proposed to mediate the effect on flowering of pepper VIL1 [22]. The identification of a global effect on H3K27me3 modification in *crel* mutants suggests that there are many more VIL targets than previously described. Together with the pleiotropic phenotypic effect, this suggests that VIL proteins are involved in a wide range of developmental processes, and play a central role in recruiting PRC2 complexes to many targets genome wide. The similarly pleiotropic effect of *Cavil1* mutants in pepper, together with its effect on gene expression [22], suggests that this is also true in other species.

### A conserved role for VIL genes in promoting flowering

VIL genes have been shown to affect flowering time in many species where mutations or silencing of these genes have been described, including both dicots and monocots [7,11,13,15,17,19,22]. In Arabidopsis, VIL proteins promote flowering by recruiting PRC2 to flowering repressors from the FLC family, thus facilitating the deposition of the repressive chromatin modification H3K27me3. Different Arabidopsis VILs act to induce flowering in specific combinations of flowering pathways, timing and target genes [8,11]. Interestingly, while tomato plants do not require vernalization for flowering and also lack *FLC, crel* mutants are late flowering. Similarly, VIL genes promote flowering in additional species that do not require vernalization for flowering and do not have a close homolog of FLC, or in species with a different vernalization mechanism. In rice, OsVIL2 and OsVIL3/LC were shown to act by repressing *OSLFL1* and *OsLF*, respectively, two flowering repressors unrelated to FLC [15,19,62]. Therefore, while the effect on flowering and possibly the molecular mechanism are conserved, the target genes differ among species [22]. It will be interesting to identify the flowering repressor that mediates this effect in tomato.

## Materials and methods

### Plant material and growth conditions

Tomato plants (*Solanum lycopersicum* cv M82) were germinated and grown in a controlled growth room or in a commercial nursery for four weeks. Then the seedlings were transferred to a greenhouse with natural daylight and 25^0^C/ 20^0^C day/night temperature, or to an open field with natural daylight and temperature. *crel-1* was isolated in this work by a mutant screen in the *e-3* background (Berger 2009, Ben Gera 2012), as described below. *crel-2 - crel-5* are from the mutant populations described by Menda et al.,[41]. The transactivation system, described previously [50,51], was used to characterize the CREL promoter and for leaf-specific expression. This system consists of driver lines and responder lines. In the driver lines, specific promoters drive the expression of the synthetic transcription factor LhG4, which does not recognize endogenous plant promoters. In the responder lines, a gene of interest or a reporter is expressed downstream of the E.coli operator, recognized by LhG4 but not endogenous plant transcription factors. A cross between a driver and a responder lines results in a plant (designated PROMOTER>>GENE) expressing the gene of interest/marker under the control of the specific promoter. *La-2, clau*, the *FIL* driver line and the *ARF10, miR160* and *EdII-GUS* responder lines have been previously described [32,37,44,48,51,63]. *35S::CREL* lines were generated in this work, as described below.

### Generation of a mutant population, screening and identification of *crel-1*

Around 750 *entire-4* (*e-4*) seeds were treated with the mutagenic substance Ethyl-Methane Sulfonate (EMS, Sigma m0880) at a concentration of 0.6% for 10 hours. Around 50 seeds underwent a control treatment without exposure to EMS. The treated seeds were sown in a commercial nursery and the seedlings (M1 generation) were transferred to a greenhouse. M1 plants were self-pollinated to increase the number of seeds per plant. M2 seeds were collected separately from each of around 650 M1 plants. M2 progeny (around 40 seeds per family) were grown in an open field, and screened frequently during the season for mutants that affect the development of the leaf, flower and fruit. *e crel-1* was identified in this screen as a recessive mutant segregating 1:3 in an M1 family. Single *crel-1* mutants were generated by a cross between *e crel-1* and wild type and identification of single *crel-1* mutants in the F2 generation. *crel-1* was then back-crossed three times to M82 for further characterization.

### Allelism tests and genetic interactions

As *crel* mutants are sterile, allelism tests were performed by crossing heterozygote siblings. Progeny of a cross between two heterozygous alleles segregated ¼ mutant progeny. Similarly, genetic interactions between *crel* and other mutants or transgenic lines were generated by crossing heterozygous *crel* siblings with the respective mutant or transgenic line.

### Identification of the *CREL* gene

An F2 mapping population was generated by crossing *crel-1/+* plant, in the M82 background to *Solanum pimplinellifolium*, and collection of F2 progeny from individual F1 plants. Initial mapping with 30 F2 plants showing the *crel-1* phenotype, and 50 mapping markers developed by Revital Bronstein, Yuval Eshed (Weizmann Institute) and Zach Lippman (CSHL) and spread along the tomato genome, identified linkage to 3 markers on chromosome 5. Fine mapping of 120 *crel-1* F2 individuals and additional markers located the gene to a region between markers zach 43.2 and jose 58.1 dcap, located between bases 43,123,344 and 58,170,500 on chromosome 5. Further mapping was hampered by an introgression of *S. pimpinellifolium* sequences in the M82 line in this region [49] We therefore used RNAseq of wild type, *crel-1* and *crel-2* plants to identify polymorphism in these alleles.

For RNAseq, shoot apices containing the SAM and the 4 youngest primordia were collected from 14-day-old M82, *crel-1* and *crel-2* plants, in which L4 (the 4^th^ leaf produced by the plant) was at the P4 stage. RNA was extracted using the RNeasy micro kit^TM^ (Qiagen), using the manufacturer’s instructions. Two biological replicates were used for M82 and *crel-2*, and one biological replicate for *crel-1*. Sequencing libraries were prepared according to the Illumina TruSeq RNA protocol and sequenced on an Illumina HiSeq2000 platform at the Genome Center of the Max Planck Institute for Plant Breeding Research. We obtained between 21,3 and 28,3 million 96-bp single-end reads per library (average of 25,8 million). Reads were aligned to the *S. lycopersicum* reference sequence v2.40 using TopHat v2.0.6 [64] with the following parameters: - max-insertion-length 12 -max-deletion-length 12 -g 1 -read-gap-length 12 -read-edit-dist 20 -read-mismatches 12 -no-coverage-search -read-realign-edit-dist 0 -segment-mismatches 3 -splice-mismatches 1. To detect polymorphisms between the *crel* mutants and wild-type M82, biological replicates from each genotype were merged. Then, duplicated reads were removed using default settings in Picard (http://broadinstitute.github.io/picard/), indels were realigned using GATK v2.2-8, and variants called in all samples simultaneously using default parameters in GATK v2.2-8 [65]. Next, we estimated the effect of each variant in annotated transcripts (ITAG 2.3) using ANNOVAR [66]. Variants in the candidate region in chromosome 5 determined by QTL analysis were evaluated manually.

### Phenotyping and imaging

#### Characterization of early leaf development and rate of leaf initiation

Plants were sown, germinated and grown in a growth chamber. Every two weeks, the number of leaves and leaf primordia were counted from six plants from each genotype. The fifth leaf (L5) was photographed by a stereoscope (Nikon SMZ1270) and its developmental stage determined. Six different plants were used for each time point.

#### Quantification of leaf complexity

Leaves 5, 7 and 9 were marked at the time of their emergence from the shoot apex. Then, the number of leaflets was counted every 7-14 days for each marked leaf. Primary, intercalary, secondary and tertiary leaflets were counted. At least 9 plants were counted for each genotype.

#### SEM characterization of flower development

Flowers from different developmental stages of each genotype were collected and their petals removed using a stereoscope, placed on a microscope stub with a carbon strip and analyzed with Hitachi TM3030 Plus SEM.

#### Phenotypic quantification and statistical analysis

For the quantification of the number of leaves to flowering, plants were grown in a greenhouse, and with the appearance of the first flower, the number of leaves formed before the flower were counted. At least 9 biological repeats, each consisting of a single plant, were quantified. The experiment was repeated twice, once with plants germinated in a commercial nursery and once with plants germinated in a growth chamber. Quantification of plant height and width was performed on 15-week-old plants grown in a greenhouse. Plant height was measured from the soil to the stem tip. For the quantification of root phenotypes, seedlings were grown hydroponically in Hoagland nutrient solution (pH 6.5), in a growth room set to a photoperiod of 12/12-h night/days, light intensity (cool-white bulbs) of ~250 μmol m^-2^ s^-1^, and 25°C. After 28, 34, 38 and 43 DAG the roots of 3 plants of each genotype were scanned and analyzed using a flatbed scanner (Epson 12000XL, Seiko Epson, Japan) and root architecture was analysed using WinRhizo software (Regent Instruments Ltd., Ontario,Canada). Statistical analysis was performed using the JMP software (SAS Institute, http://www.sas.com). Means and p values were calculated using the Student’s t-test or the Dunnett’s test, as indicated in the figures.

#### Histological characterization of stem tissues

Ten to 50 day-old plants were freehand dissected using a double-sided razor blade. 1-2-mm-long sections were dissected from up to 5 cm below the node. Sections were dehydrated in acetic acid: ethanol [1:10] for 1 hour and then stained directly with TBO (0.01% aqueous, sigma). Images of early developmental stages were captured using Nikon a SMZ1270 stereoscope equipped with a Nikon DS-Ri2 camera and NIS-ELEMENTS software.

#### Confocal imaging

For analysis of pCREL:nYFP expression, dissected whole-leaf primordia were placed into drops of water on glass microscope slides and covered with cover slips. The pattern of YFP expression was detected by a confocal laser scanning microscope (CLSMmodel SP8; Leica), with the solid-state laser set at 514 nm for excitation and 530 nm for emission. Chlorophyll expression was detected at 488nm for excitation and 700nm for emission. ImageJ software was used for analysis, quantification, and measurements of captured images. Images were adjusted uniformly using Adobe Photoshop CS6. Tomato stems and primary roots were cut to sections of 200um and 300um width respectively using Leica VT1000 vibratome and were and cleared using ClearSee [67], cell wall staining was performed using SR2200 (Renaissance Chemicals) prior to mounting and visualization using 405nm laser.

### Cloning and plant transformation

The *CREL* promoter was generated by amplifying 3000 bp upstream of the *CREL* ATG from genomic DNA and cloned upstream to LhG4 generating the *pCREL:LhG4* driver line.

The *CREL* gene was amplified from tomato M82 cDNA using the op:VIN3 F and op:VIN3 R primers (S2 Table), and cloned into the pENTR/d™ vector using a TOPO isomerase cloning system (Invitrogen). The *CREL* gene was then subcloned using L/R clonase (Invitrogen) downstream to the 35S promoter to generate *35S::CREL*.

Plant transformation and tissue culture were performed as described in Israeli et al 2019 [36]. At least five independent kanamycin-resistant transgenic lines from each transgene were genotyped and, in the case of *pCREL:LhG4*, crossed to an *OP:YFP* stable line to generate pCREL>>YFP. Three lines from each transgene or resultant cross were examined, and a representative line was selected for further analysis.

### Phylogenetic Analysis

Phylogenetic analysis was performed using full-length protein sequences of the tomato, Arabidopsis, rice and pepper VIL gene family. The sequences were obtained from the Sol Genomics Network (SOL, https://solgenomics.net/), The Arabidopsis Information Resource (TAIR, https://www.arabidopsis.org/) and the Plaza tool (https://bioinformatics.psb.ugent.be/plaza/). Sequences were aligned and phylogenetic tree was constructed using Clustal W (https://www.ebi.ac.uk/Tools/msa/clustalo/) [68,69].

### RNA extraction and qRT-PCR analysis

RNA was extracted using the Plant/Fungi Total RNA Purification Kit (Norgen Biotek, Thorold, ON, Canada) according to the manufacturer’s instructions, including DNase treatment. cDNA synthesis was performed using the Verso cDNA Kit (Thermo Scientific, Waltham, MA, USA) using 1 μg of RNA. qRT-PCR analysis was carried out using a Corbett Rotor-Gene 6000 real-time PCR machine, with SYBR Premix for all other genes. Levels of mRNA were calculated relative to *EXPRESSED* (*EXP*) [70] or *TUBULIN* (*TUB*) [71] as described [26]. Primers used for the qRT-PCR analysis are detailed in S2 table.

### ChIP-seq procedures

WT and *crel* plants were grown on soil under 16 hrs of light/8 hrs dark cycles for 28 days after germination. Shoot apices, (0.8 g for each replicate and two replicates per genotype) containing approximately three visible expanding leaves, were harvested and fixed in 1% formaldehyde + 0.2% Silwet L-77 for 17 minutes under vacuum. Glycine was then added to a final concentration of 0.125 M and tissue was placed under vacuum for an additional 5 minutes, followed by washing several times in water. ChIP was performed on the fixed tissue using the procedure of Gendrel et al. [72]. For each ChIP reaction, we used 2 ug of a rabbit polyclonal antibody against H3K27me3 (Millipore, catalog #07-449). Input and ChIP DNAs were converted to Illumina sequencing libraries using the Accel-NGS 2S Plus DNA library kit according to the manufacturer’s instructions (Swift Biosciences). Libraries were sequenced on an Illumina NextSeq 500 instrument using 50-nt single end reads at the University of Georgia Genomics and Bioinformatics Facility.

### ChIP-seq data processing

Raw reads were mapped to the SL3.0 build of the tomato genome using Bowtie2 [73] with default parameters. Raw mapped reads were then processed using Samtools [74] to retain only those with a mapping quality score greater than or equal to 2. Enriched regions (peaks) for H3K27me3 were then identified for each replicate using the “*Findpeaks*” function of the HOMER package [75]. Further analyses only considered peaks that were identified in both replicates for each genotype. For normalization and visualization, quality-filtered reads in *.bam* format were converted to *bigwig* format using the “*bamcoverage*” script in deepTools 2.0 [76] with a bin size of 1 bp and RPKM normalization. Heat maps and average plots displaying ChIP-seq data were also generated using the “*computeMatrix*” and “*plotHeatmap*” functions in the deepTools 2.0 package.

## Data accessibility

RNAseq reads are available at https://www.ncbi.nlm.nih.gov/sra under project numbers PRJNA347502 (M82) and PRJNA723668 (crel-2 and crel-2).

All ChIP-seq datasets have been deposited to the NCBI GEO database and are available under accession number GSE174416.

## Acknowledgments

We thank Yuval Eshed, Alon Samach and Dani Zamir for plant material and fruitful discussions and suggestions; members of the Ori lab for continuous discussions and help; Revital Bronstein, Yuval Eshed and Zach Lippman for sharing the tomato mapping marker set.

## Supporting information captions

**S1 Fig. Characterization of *crel* mutants.** (A-D) Mature 5^th^ leaves of the indicated genotypes.White arrowheads point to primary and intercalary leaflets. Scale bars: 2cm. (E) Slower leaf production in *crel-2*. The Y axes shows the developmental stage (plastochron, P) of the fifth leaf produced by the plant at the indicated days after seeding. Error bars indicate SD (n=5-11).

**S2 Fig. *35S:CREL* has sublte developmental phenotype alterations when compared with WT.** (A) qRT-PCR analyzing CREL expression levels in *35S:CREL* shoot apices containing the SAM and 5-6 young leaf primordia. Each repetition contained 9 or more plants. Error bars indicate SE (n = 3). Asterisks indicate statistically significant differences by student t-test, *P < 0.05. (B) Plant hight, measured from the cotyledons to the tip of the plants at the end of the growing season, on 15-week-old plants. Error bars represent the SE of 5 (*crel-2*), 3 (wt) or 5 (*35S:CREL*) repeats; *p*-values indicate differences from WT, as determent by Dunnett’s test. (C) Flowering time of the 3 independent *35S:CREL* lines in comparison to the wild type, measured by number of leaves produced before flowering. Error bars represent the SE of 5-12 plants; *p*-values indicate differences from WT, as determent by Dunnett’s test. (D) Total number of leaflets, measured on expanded 5^th^ leaves. Error bars represent the SE of 12 plants for the wild type and 6 plants for each of the *35S:CREL* lines. *p*-values indicate differences from WT, as determent by Dunnett’s test.

**S3 Fig. Overview of ChIP-seq datasets.** Principal component analysis of input DNA and ChIP-seq samples.

**S1 Table. Sequence read numbers for ChIP-seq.** Two biological replicates of ChIP-seq for H3K27me3 were performed on shoot apices of WT and *crel1-2* plants. The table indicates for each biological replicate the number of total sequencing reads obtained, the number and percentage mapping to the tomato genome, and the total number of reads remaining after filtering for mapping quality. Reads with a MAPQ score of 2 or greater were used for further analysis.

**S2 Table. Primers used in this work**.

